# Transcriptomic profiling of nematode parasites surviving after vaccine exposure

**DOI:** 10.1101/144980

**Authors:** Guillaume Sallé, Roz Laing, James A. Cotton, Kirsty Maitland, Axel Martinelli, Nancy Holroyd, Alan Tracey, Matthew Berriman, W. David Smith, George F. J. Newlands, Eve Hanks, Eileen Devaney, Collette Britton

## Abstract

Some nematode species are economically important parasites of livestock, while others are important human pathogens causing some of the most important neglected tropical diseases. In both humans and animals, anthelmintic drug administration is the main control strategy, but the emergence of drug-resistant worms has stimulated the development of alternative control approaches. Among these, vaccination is considered to be a sustainable and cost effective strategy. Currently, Barbervax^®^ for the ruminant strongylid *Haemonchus contortus* is the only registered subunit vaccine for a nematode parasite, although a vaccine for the human hookworm *Necator americanus* is undergoing clinical trials (HOOKVAC consortium). As both these vaccines comprise a limited number of proteins there is potential for selection of nematodes with altered sequence or expression of the vaccine antigens. Here we compared the transcriptome of *H. contortus* populations from sheep vaccinated with Barbervax^®^ with worms from control animals. Barbervax^®^ antigens are native integral membrane proteins isolated from the brush border of the intestinal cells of the adult parasite and many of them are proteases. Our findings provide no evidence for changes in expression of genes encoding Barbervax^®^ antigens in the surviving parasite populations. However, surviving parasites from vaccinated animals showed increased expression of other proteases and regulators of lysosome trafficking, and displayed up-regulated lipid storage and defecation abilities that may have circumvented the vaccine effect. Implications for other potential vaccines for human and veterinary nematodes are discussed.

## 1. Introduction

Gastrointestinal nematodes (GIN) are clinically and economically important parasites of humans (Hotez et al., 2016) and livestock species (Kaplan and Vidyashankar, 2012). Human GIN (e.g. hookworm, roundworm and whipworm) infect over one billion people worldwide, resulting in the loss of over three million disability-adjusted life years (DALYs) in 2013 (Hotez et al., 2016). In ruminants, parasitic nematode infections cost the global livestock industry billions of dollars annually in production losses and treatments (Kaplan and Vidyashankar, 2012). GIN therefore impede both human health and wealth and are an aggravating factor of poverty (Rist et al., 2015).

Control of veterinary parasites has relied primarily on strategic drug administration (McKellar and Jackson, 2004). However the increase in anthelmintic resistance, particularly multidrug resistance, threatens the viability of the livestock industry in many regions of the world (Kaplan and Vidyashankar, 2012). Similarly, control of human helminthiases involves large-scale community treatment, which has resulted in reductions in GIN prevalence over the last 30 years (Hotez et al., 2016; Clarke et al., 2017). However, there is also the potential for anthelminthic drug failure (Hotez et al., 2016) and recent surveys have indicated a low or variable cure rate of human hookworm infection after benzimidazole treatment, with no reduction in anaemia in some endemic regions (Keiser and Utzinger, 2008; Soukhathammavong et al., 2012).

It is unlikely that novel anthelmintic compounds will be approved at an equivalent pace to the emergence of anthelmintic resistance (Geary et al., 2004). Greater research effort is therefore being directed at vaccine development for more sustainable GIN control in both veterinary and human settings (Hewitson and Maizels, 2014; Hotez et al., 2016). Vaccines may be used alone or combined with drug treatment to reduce the emergence of drug resistance (Lee et al., 2011). In comparison to antimicrobial drugs, there are few examples of the development of resistance to vaccination in bacterial or viral pathogens (Kennedy and Read, 2017). However, the antigenic complexity and immunoregulatory capacity of nematode parasites makes vaccine development challenging (Hewitson and Maizels, 2014). Only two vaccines are currently commercially available: Barbervax^®^ licensed in Australia in 2014 and comprising native parasite gut membrane glycoproteins of the ovine GIN *Haemonchus contortus* (Bassetto and Amarante, 2015; Kearney et al., 2016), and Bovilishuskvac^®^, an irradiated larval vaccine for the cattle lungworm *Dictyocaulus viviparus* (McKeand, 2000). For the human hookworm *Necator americanus*, a phase 1 clinical vaccine trial has been carried out using recombinant aspartic protease Na-APR-1 combined with gluthatione-S-transferase-1 (Na-GST-1) (Brelsford et al., 2017).

Digestion of haemoglobin in haematophagous nematodes like *H. contortus* or hookworms requires activity of different proteolytic enzymes, including aspartic, cysteine and metallo-proteases and exopeptidases (Williamson et al., 2003) underscoring the large expansion of protease gene families identified within the genome of *H. contortus* (Laing et al., 2013; Schwarz et al., 2013). Barbervax^®^ is prepared from gut membrane extracts of *H. contortus* adult worms and contains two major protease fractions, H11 and H-gal-GP (Smith et al., 2001). H11 is a family of microsomal aminopeptidases for which five isoforms have been identified in native extracts (Munn et al., 1997; Roberts et al., 2013), and several related isoforms recently found from genome and transcriptome analysis (Mohandas et al., 2016). H-gal-GP is a 1,000 kDa complex of four zinc metallopeptidases (MEP1-4) and two pepsinogen-like aspartyl proteases (PEP-1 and PEP-2) (Smith et al., 2003), together with additional components (thrombospondin, galectins and cystatin), thought unlikely to be protective (Knox et al., 2003). Vaccination of sheep with either H11 or H-gal-GP individually reduced worm burden and faecal egg count by 70% and 95%, respectively (Munn et al., 1997; Newton and Munn, 1999; Knox et al., 2003; LeJambre et al., 2008; Roberts et al., 2013). Cysteine proteases HmCP-1,4 and 6, enriched from adult *H. contortus* gut membrane provided a lower level of protection (Knox et al., 2005). Barbervax^®^ induces circulating antibodies which are ingested by the parasite when it feeds and which inhibit haemoglobinase activity *in vitro* (Ekoja and Smith, 2010) and probably *in vivo*. Because the gut-membrane antigens are not exposed to the host immune system during natural infection, Barbervax^®^ relies on the induction of antibodies to “hidden” antigens (Knox et al., 2003). Therefore, it is speculated that the Barbervax^®^ proteins are not under selective pressure during natural infection, but whether vaccine-induced immunity influences levels of gene expression is currently unknown.

The high level of genetic diversity observed in genomic datasets of *H. contortus* (Laing et al., 2013) and other helminths underpins their capacity for adaptation and contributes to the evolution of drug resistance (Gilleard and Redman, 2016). It is clear that pathogens can evolve in response to other interventions, including vaccination, in some cases leading to vaccine escape and failure (Brueggemann et al., 2007; Kennedy and Read, 2017). Given the limited number of antigens composing the *H. contortus* vaccine, selection may arise in the field. Here we compare the transcriptomes of *Haemonchus* adults surviving in Barbervax^®^ vaccinated animals with worms recovered from control animals post challenge infection. Identifying any effects that vaccines may have on helminth populations may guide their optimal use in the field.

## 2. Materials and methods

### 2.1 Experimental design and collection of parasite material

Adult worms examined in this study were collected on completion of a Barbervax^®^ vaccine trial carried out at the Moredun Research Institute, UK. Twelve six month old worm-free Texel cross lambs were allocated into groups of six, balanced for sex and weight. One group was injected subcutaneously with two doses of Barbervax^®^ four weeks apart, whilst the second, control group was not vaccinated. All sheep were given a challenge infection of 5,000 *H. contortus* MHco3(ISE) L3 administered *per os* on the same day as the second vaccination. The MHco3(ISE) strain is susceptible to all broad-spectrum anthelmintics (Roos et al., 2004) and was inbred to produce the material for the *H.contortus* genome sequencing project at the Wellcome Trust Sanger Institute (Laing et al., 2013). All strains were maintained at the Moredun Research Institute. The same *H. contortus* MHco3(ISE) strain was used to generate the vaccine for this study and to challenge vaccinated and control lambs.

Fecal egg counts (FEC) were monitored twice weekly between days 17 and 29 post-challenge by a McMaster technique with a sensitivity of 50 eggs/g. Adult worms were recovered from each sheep at post-mortem 31 days post-challenge. Antibody titres were measured by ELISA, with plates coated with Barbervax^®^ (50 μl per well at 2 μg/ml). Serum samples were serially diluted (from 1/100 to 1/51200) in PBS/0.5% Tween and binding detected using mouse anti-sheep IgG (Clone GT-34, Sigma G2904; 1:2500 dilution) and rabbit anti-mouse IgG-HRP conjugate (Dako P0260; 1:1000 dilution). Antibody titres are expressed as the reciprocal of the end-point dilution resulting in an OD of ≥ 0.1 above the average negative control value.

### 2.2 Ethics Statement

Experimental infections were performed at the Moredun Research Institute, UK as described previously (Laing et al., 2013). All experimental procedures were examined and approved by the Moredun Research Institute Experiments and Ethics Committee (MRI E46 11) and were conducted under approved UK Home Office licence (PPL 60/03899) in accordance with the 1986 Animals (Scientific Procedures) Act.

### 2.3 Extraction protocol, library preparation and sequencing

To avoid any confounding factors from eggs in females or differences in sex ratio between samples, only male worms were used for RNA sequencing. RNA sequencing was carried out on pools of seven to ten surviving *H. contortus* adult worms from each animal. In total, 54 worms that survived following challenge infection of the Barbervax vaccinated sheep (V group) and 60 worms from control sheep (C group) were picked for RNA preparations (supplementary table S1).

Total RNA was extracted from the worms using a standard Trizol (Thermo Fisher Scientific, 15596026) protocol and libraries prepared with the Illumina TruSeq RNA preparation kit before sequencing using a HiSeq 2500 platform with v3 chemistry.

### 2.4 Real-time PCR

Total RNA was extracted from triplicate samples of five female worms from the same populations as the sequenced males. 3 μg total RNA was used per oligo(dT) cDNA synthesis (SuperScript^®^ III First-Strand Synthesis System, ThermoFisher, 18080051) with no-reverse transcriptase controls included for each sample. cDNA was diluted 1:100 for quantitative RT-PCR (RT-qPCR) and 1ul added to each reaction. RT-qPCR was carried out following the Brilliant III Ultra Fast SYBR QPCR Master Mix protocol (Agilent Technologies, 600882) and results analysed using MxPro qPCR Software, Version 4.10. Gene expression was normalised to *ama* (HCOI01464300) and *gpd* (HCOI01760600) (Lecova et al., 2015). Primer sequences are listed in Table S2.

### 2.5 Improved *H. contortus* assembly and corresponding gene model

The *H. contortus* MHco3.ISE reference genome assembly used for this study was a snapshot of the latest version as of 14/11/2014. This assembly consists of 6,668 scaffolds with a total assembly length of 332,877,166 bp; of which 22,769,937 bp are sequence gaps. The N50 scaffold length is 5,236,391 bp and N90 length is 30,845 bp. Specifically for this project, preliminary gene models were annotated on this assembly by transferring the gene models from the published (v1.0) genome assembly (Laing et al., 2013) using RATT (Otto et al., 2011) with default parameters, and with a *de novo* approach using Augustus v2.6.1 (Stanke et al., 2004) with exon boundary ‘hints’ from the RNAseq data described previously (Laing et al., 2013), mapped against the new reference genome in the same way as in this previous paper.

### 2.6 RNAseq data handling and differential expression analysis

RNAseq data were mapped onto the reference genome using a gene index built with Bowtie2 (Langmead and Salzberg, 2012) and TopHat v2.1.0 (Trapnell et al., 2009) with maximal intron length of 50 Kbp and an inner mate distance of 30 bp that identified 48.8% of the reads being mapped unambiguously to a gene feature. Counts of reads spanning annotated gene features were subsequently determined with HTSeq v0.6.0 (Anders et al., 2015).

To ensure our biological conclusions are not sensitive to details of the statistical methods used, we implemented two different analysis frameworks for the RNA-seq count data, using the DESeq2 v1.12.4 framework (Love et al., 2014) and the *voom* function as implemented in the LIMMA package v3.28.21 (Law et al., 2014) in R v3.3.1 (R Core Team, 2016). Genes found to be significantly differentially expressed (DE, adjusted p-value <5%) by both VOOM and DESeq2 analyses were retained. A gene ontology enrichment analysis was performed using the TopGO package v2.26.0 (Alexa, 2016).

Gene identifiers of the vaccine core components, namely MEP-3 (Smith et al., 2000), MEP-1,2,4, PEP-1 (Britton et al., 1999) and PEP-2 (Smith et al., 2003) as well as H11, were retrieved via a BLAST search of their nucleotide sequence against the *H. contortus* MHco3.ISE reference assembly (Laing et al., 2013) in WormBase ParaSite (Howe et al., 2016). The expression levels of candidate housekeeping genes (Lecova et al., 2015) were also retrieved using the gene identifiers associated with their GenBank records (Table 1).

**Table 1.**
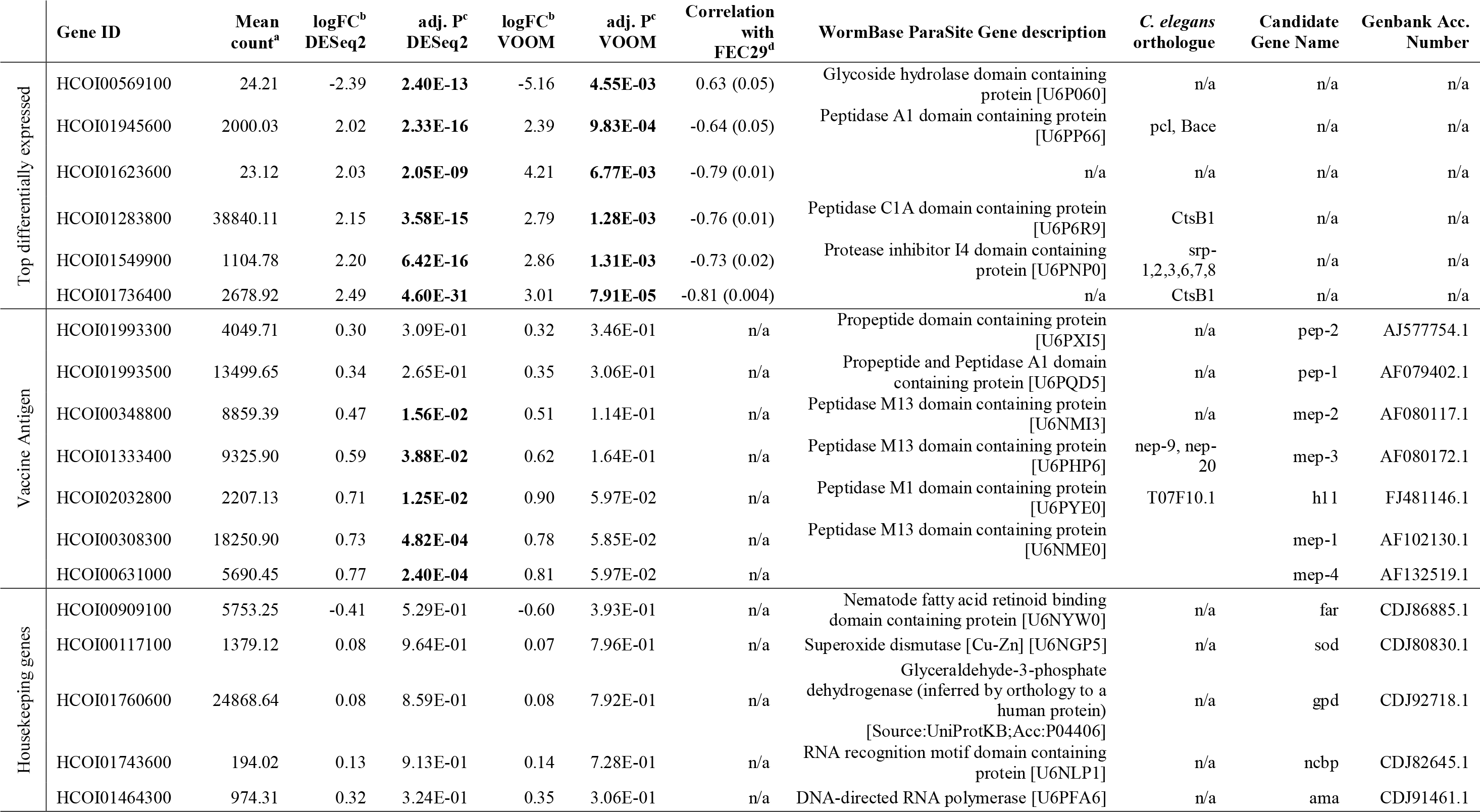
Gene of interest expression levels, fold change and associated p-values.

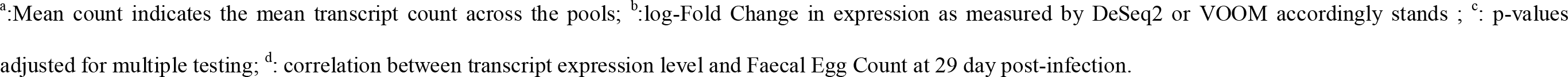

## 3. Results

### 3.1. Vaccination greatly reduces faecal egg counts in vaccinated sheep

Parasitological data confirmed a significant reduction in *H. contortus* infection following Barbervax vaccination. Over the course of the trial, vaccinated sheep (Group V) shed significantly fewer eggs (mean 390 ± 639 eggs per gram faeces (epg), Fig 1A, Table S1) than the control group (Group C) given the same challenge infection dose without prior vaccination (mean 5,914± 2,628 epg), representing a 15-fold decrease (Wilcoxon test, p=0.002). Vaccinated sheep contained fewer worms, indicated by the significantly lower worm volume collected at necropsy compared to control sheep (2.8 mL ± 1.9 versus 6.7 mL ± 3.5; Table S1). Among the V group, V_5 showed an outlying egg excretion over the course of the trial (1,647 epg at necropsy; upper 95% confidence interval limit of 861 epg estimated after 1,000 bootstraps), suggesting a relatively suboptimal vaccine response in this animal. This is supported by the lower antibody titre of this sheep, relative to the other Barbervax vaccinated animals, at day 28 post-challenge infection (Fig 1B).

**Figure 1.**
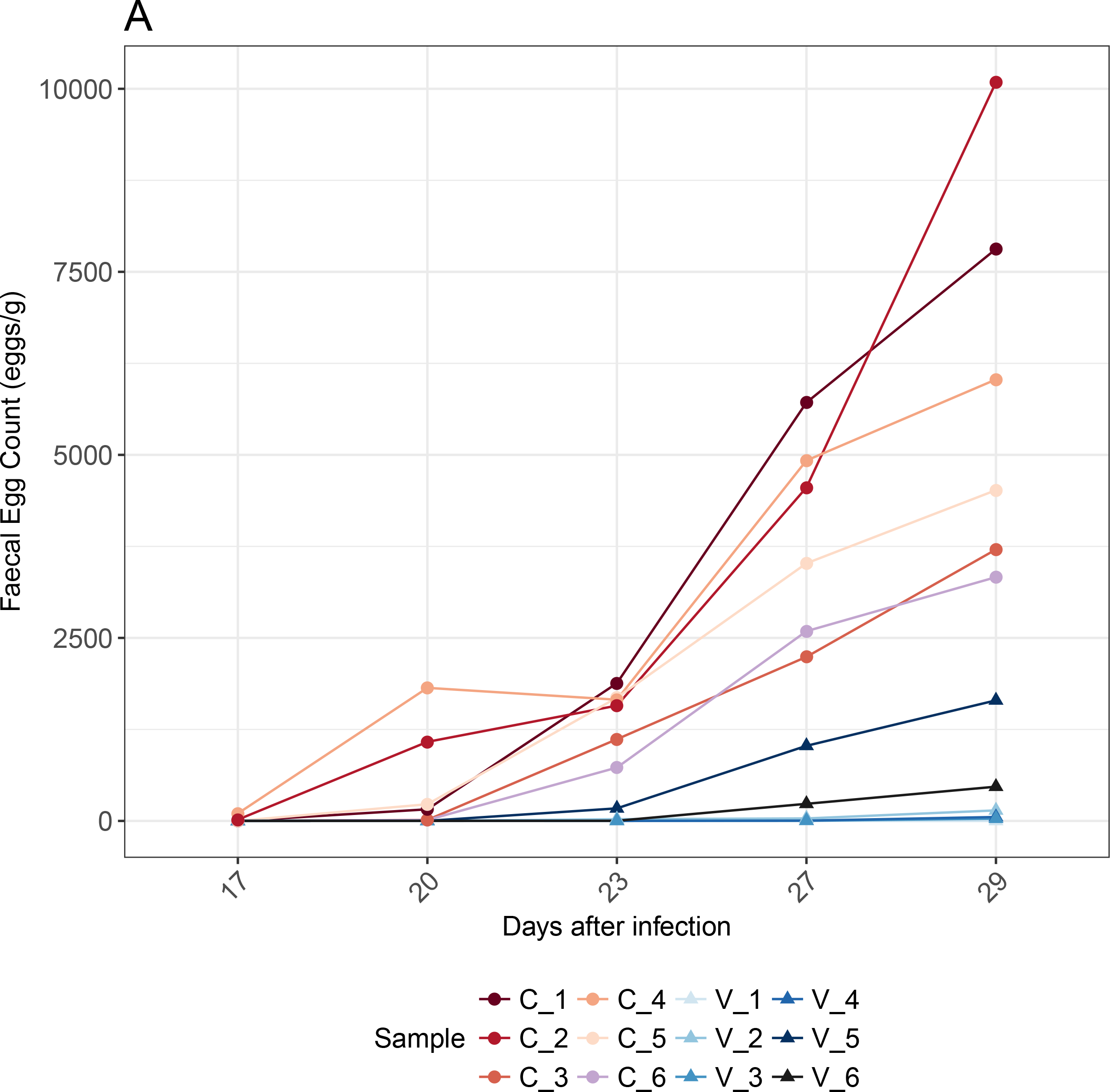
Faecal egg counts and anti-Barbervax IgG titer of individual sheep. Fig 1A Faecal egg counts from each of the 12 sheep in the trial were plotted for each available time point post challenge. The plot shows a 15-fold difference in egg excretion between vaccinated and control sheep on day 29 post challenge infection. Dots for V_1, V_3 and V_4 overlap around 0 as a result of low counts. Fig 1B Faecal egg count measured at necropsy plotted against respective anti-Barbervax^®^ vaccine IgG titer, showing a negative correlation between vaccine response and egg count.

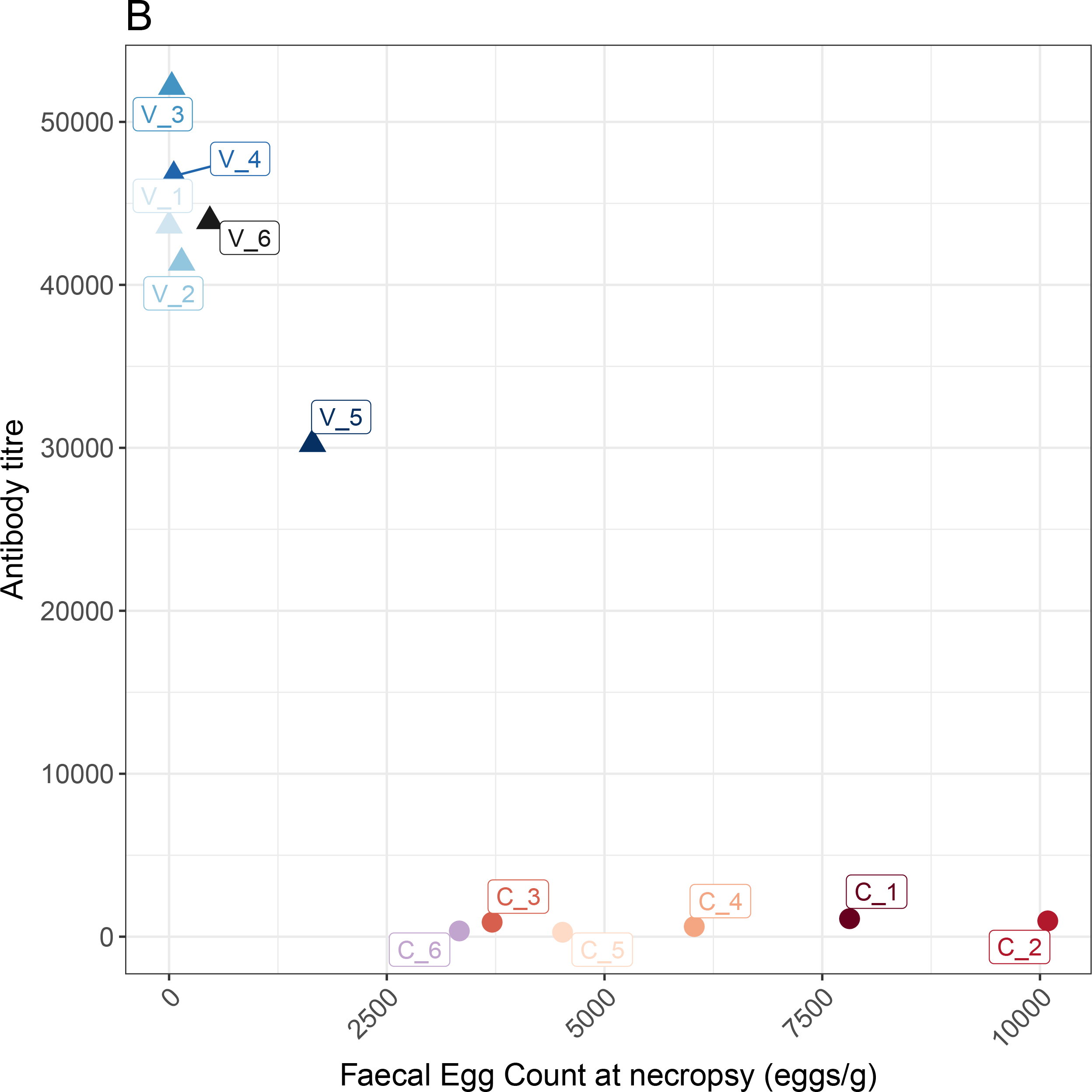

### 3.2. Transcriptional response of worms to host vaccination is dominated by higher expression of proteases and protease inhibitors

We investigated any changes in *H. contortus* gene expression in worms surviving in vaccinated sheep relative to those surviving in controls. On average 11M (standard deviation of 1.79M) reads were available for each library (Table S1). In PCA of the normalized RNA-seq read counts, the first two axes explained 53% of the total variation, 37% of which was resolved along the 1^st^ axis that separated the experimental groups (Fig S1). Two pools of worms sampled from control sheep, C_4 and C_6, showed atypical behaviour that was resolved along the 2^nd^ PCA axis (Fig S1). These samples were discarded from the dataset for subsequent analyses, resulting in a comparison of 6 V samples and 4 C samples.

We found 52 genes significantly differentially expressed (DE; adjusted p-value < 0.05) between the two experimental groups, with six genes exhibiting a fold change above 4, and 34 genes showing a fold change above 2 (Figures 2 and S2, Table 1 and S3). Adult worm survival following vaccination was associated with an increase in expression of most of the DE genes, *i.e.* 46 out of 52. Among the top six DE genes, the only down-regulated gene was a glycoside hydrolase domain-containing protein (HC0I00569100, Table 1, Figure 2A). Three of the most highly up-regulated genes encoded proteins containing peptidase domains (HCOI01945600, HCOI01283800, Table 1, Figure 2A), or a peptidase inhibitor I4 domain (HCOI01549900, Table 1, Figure 2A), while two genes were unannotated (HCOI01623600, HCOI01736400). Noticeably, orthologs of HCOI01736400 in *D. viviparus* (nDv.1.0.1.g04423) or *A. caninum* (ANCCAN_06626 and ANCCAN_06627) also encoded cathepsin B (cysteine peptidase). Expression of the peptidases (HCOI01945600, HCOI01283800) and HCOI01736400 was validated by quantitative RT-PCR in female worms from the same population as the sequenced males, and confirmed a two to three-fold greater expression of each mRNA in worms surviving in vaccinated sheep compared to controls (Figure 2B).

**Figure 2.**
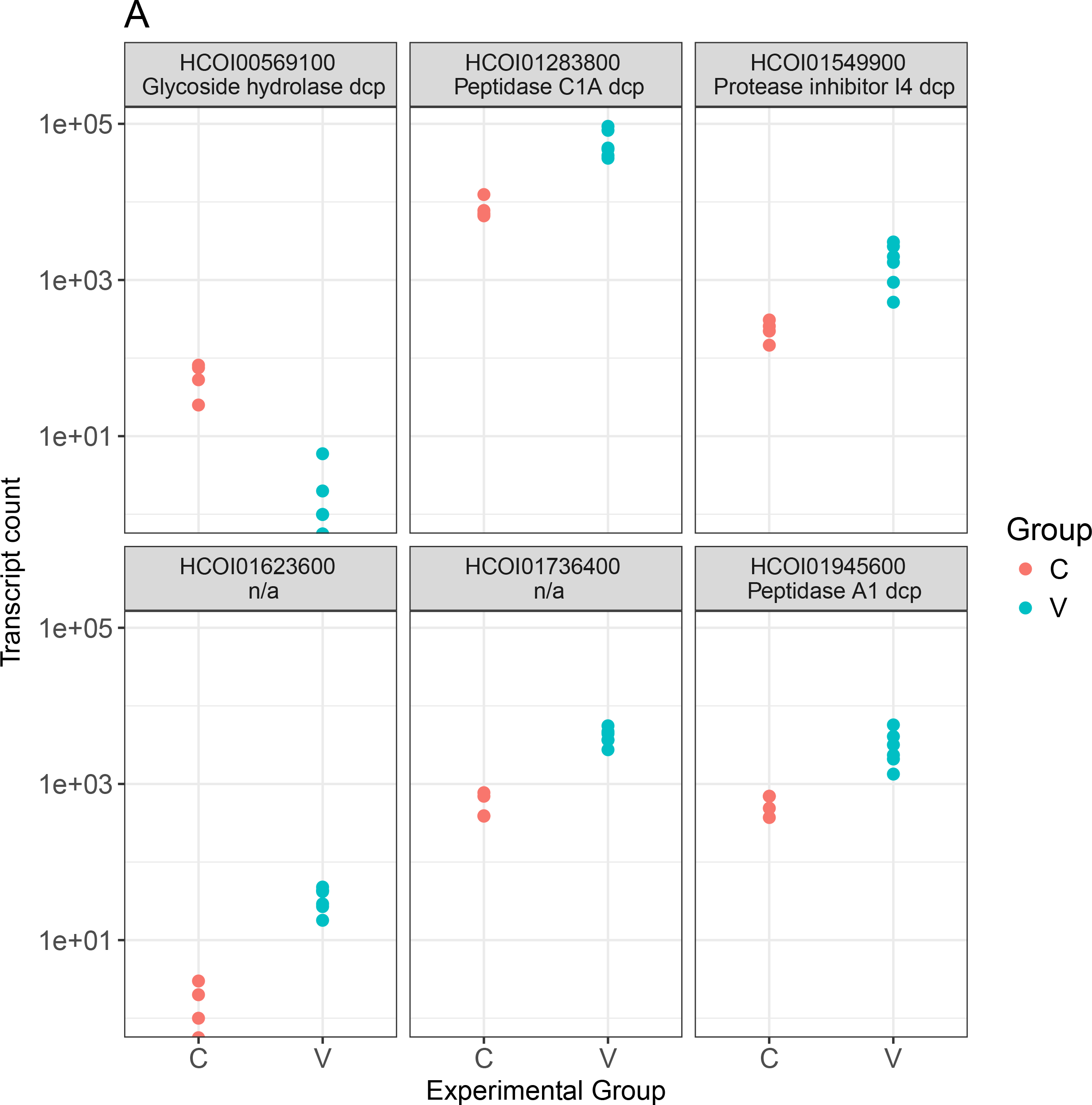
Expression level of the top differentially expressed genes within each experimental group (2A) and associated correlation with faecal egg count in control populations (2B) A. A boxplot for all six genes that exhibited an absolute log-transformed fold change of 2 between the experimental conditions. Dcp stands for “Domain Containing Protein”. B. Fold change in expression level of selected genes, by qRT-PCR, shown relative to C control population. qRT-PCR was carried out on RNA extracted from adult female worms. HCOI01283800: Peptidase C1A domain containing protein; HCOI01549900: Protease inhibitor I4 domain containing protein; HCOI01736400: ortholog to cathepsin B in *D. viviparus* and *A. caninum*.

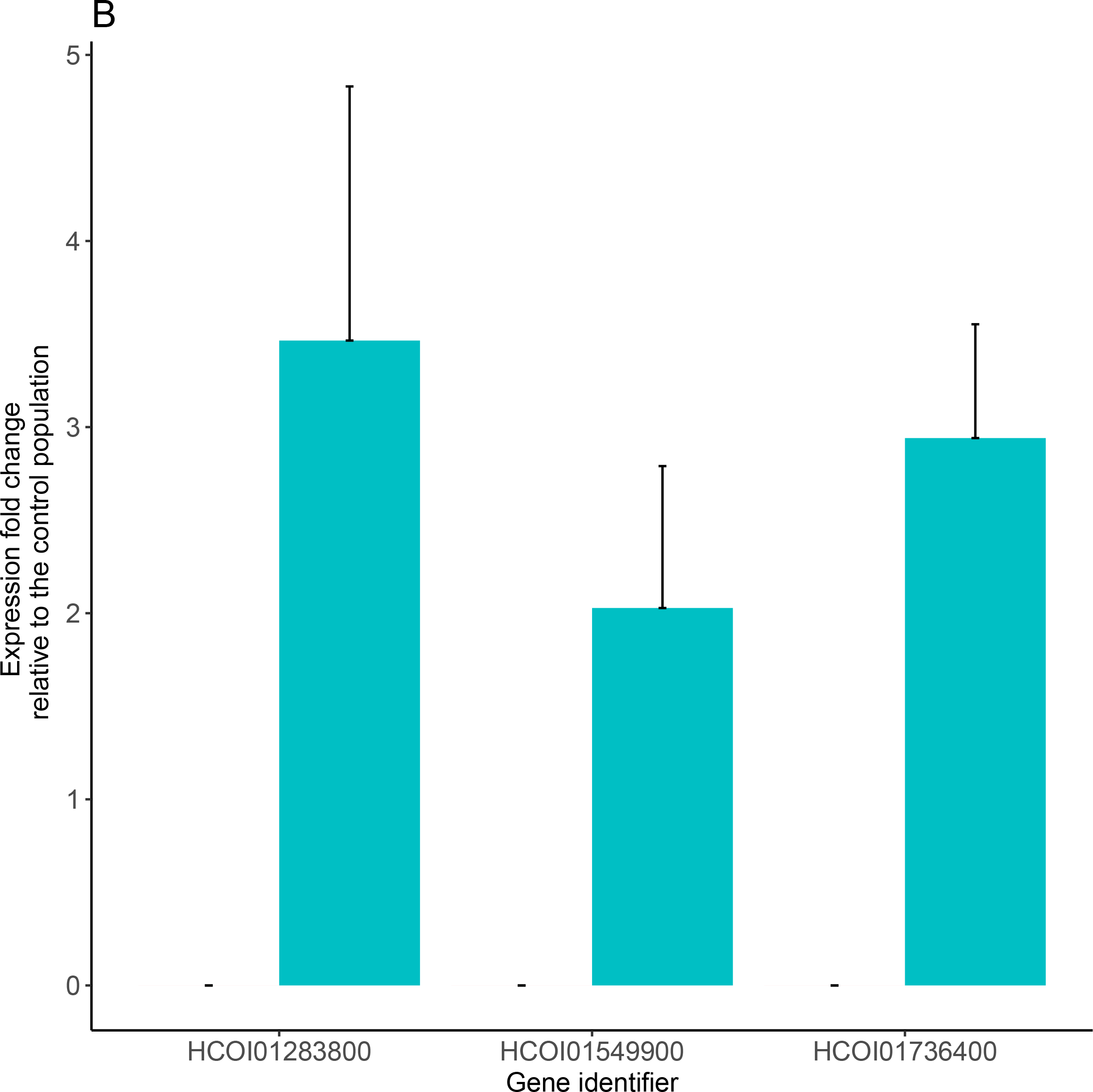

Most of the top six DE genes generally exhibited low transcript counts in control C populations (Table S4), suggesting that their higher expression in worms from group V may be triggered or selected for by the vaccine exposure. Interestingly, 14 genes among the 52 DE gene set encoded peptidases or peptidase inhibitors exemplified by the significant enrichment for peptidase activity (*p*=6.7 × 10^−15^), serine-type (*p*=9.6 × 10^−8^) and cysteine-type peptidase (*p*=2.8 × 10^−10^) GO terms (Table S5). This shift toward peptidase activity is also consistent with down-regulation of the gamma interferon-inducible lysosomal thiol reductase (*GILT*, HCOI02049600, Table S3), which is known to catalyse the reduction of cysteine proteases.

Higher expression of two genes involved in the anti-microbial response, the Lys-8 encoding gene (HCOI00041100) associated with lysozyme formation, and the anti-microbial peptide theromacin coding gene (HCOI00456500), was also found in worms surviving in vaccinated animals. A proteinase inhibitor (HCOI01591500) and a prolyl-carboxypeptidase encoding gene (HCOI01624100) showing 99.6% similarity with contortin 2 (Genbank CAM84574.1, BLASTP, e-value=0) also showed significantly greater expression in the V group (Table S3).

### 3.3. Vaccine antigen coding genes are not differentially expressed between experimental groups

Importantly we found that most of the genes encoding the core components of the Barbervax^®^ vaccine (MEPs, PEPs, Aminopeptidases) were not significantly differentially expressed between V and C worms or where significant, showed slight over-expression in the V worm population (Table 1, Figure 3).

**Figure 3.**
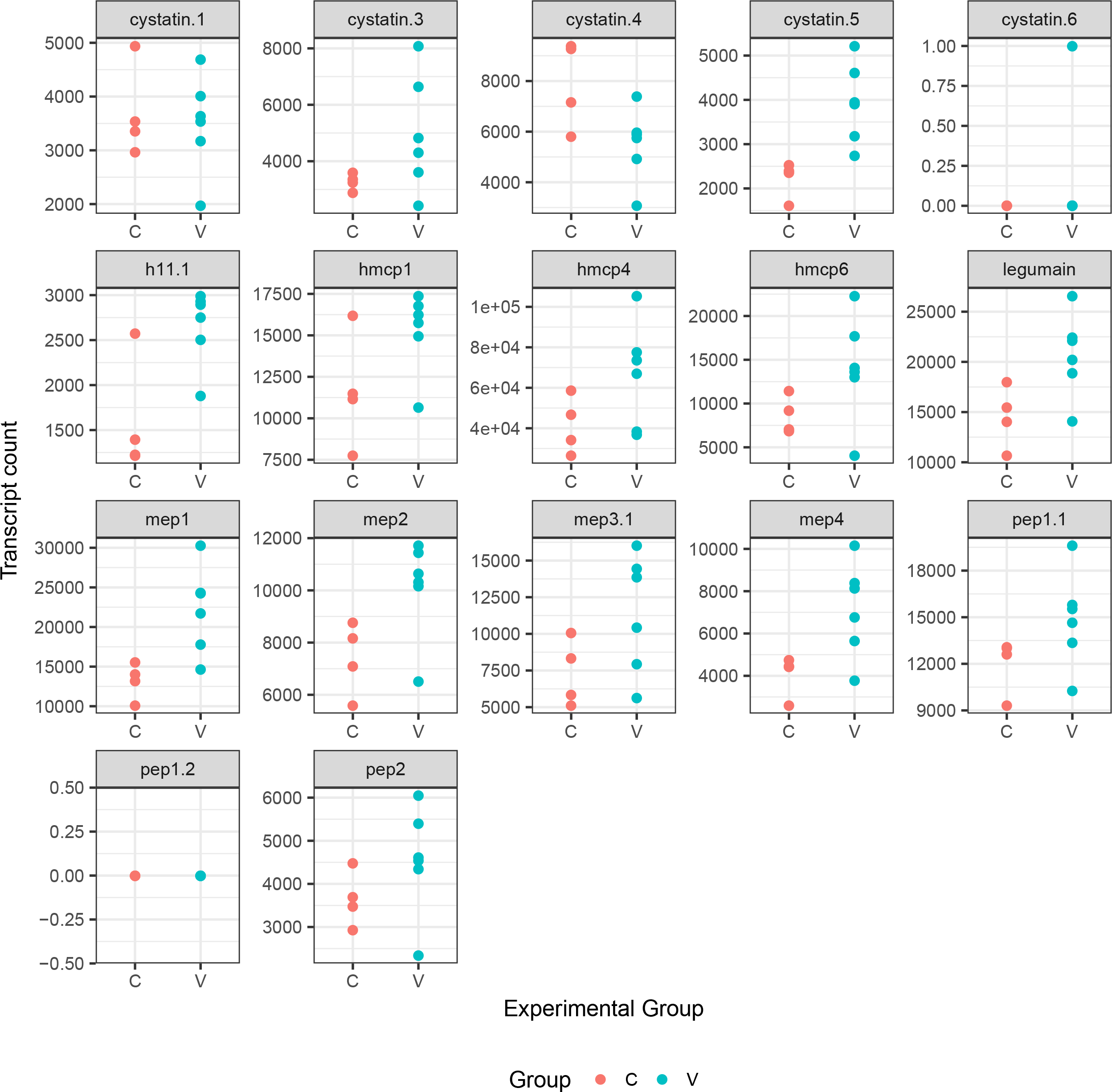
Expression level for the vaccine antigen coding genes. Figure 3 shows the normalized transcript counts for known vaccine antigen coding genes. Each dot stands for the transcript count measured in a pool of worms from vaccinated (V, green dots) or control (C, red dots) sheep. Some of the dots overlap because of similar expression levels.

## 4. Discussion

In comparison to the development of drug resistance, vaccine resistance has rarely been reported in viruses or bacteria (Kennedy and Read, 2017). These contrasting findings may relate both to the prophylactic use of vaccines, which prevent the spread of resistant mutants among hosts, and the multiplicity of pathways targeted by the host immune response following vaccination (Kennedy and Read, 2017). However, highly diverse populations, such as *H. contortus* (Gilleard and Redman, 2016) likely encompass a wide range of genotypes that could be differentially selected, ultimately leading to vaccine resistance through replacement (Martcheva et al., 2008; Weinberger et al., 2011; Barnett et al., 2015).

Resistance to all but the newest anthelmintic drugs is common and widespread amongst gastrointestinal nematode parasites of ruminants. Barbervax^®^, which is specific for *H. contortus*, is the only vaccine registered for a gut dwelling nematode of any host. While this vaccine provides a useful level of protection mediated mainly by reducing pasture contamination, a small proportion of worms do survive vaccination. Here, we investigated whether the transcriptome of these survivors differed from those of control worms.

In order to generate enough genetic material for sequencing and to avoid any contamination by egg-specific transcripts, this study focused on male worms only. Consequently, our experiment could not resolve the observed sex-specific effect of the Barbervax^®^ vaccine, *i.e.* the vaccine being more efficient on females than males (Smith and Smith, 1993), although we were able to confirm some of the observed transcriptional differences in female worms recovered from the same animals. Our data shed light on transcription modifications involved in the survival of male worms and provided insights into the mechanisms associated with their survival following vaccination.

Since both experimental groups exhibited similar levels of vaccine antigen transcripts, there was no evidence for increased expression of vaccine targets which could mediate vaccine survival. However a metallopeptidase and an exopeptidase, belonging to the same functional families (Rawlings et al., 2010) as the vaccine MEP (M13 peptidase) and H11 (M1 peptidase) respectively, were over-expressed in the vaccine survivors although it is not clear whether these could compensate for vaccine peptidases. Instead, survival following Barbervax^®^ vaccination was associated with enhanced expression of a limited subset of genes, mainly encoding cysteine peptidases. Differential tuning of a GILT-like gene, *i.e.* down-regulated in worms surviving the vaccine response, would also support proteolytic function as an important feature for vaccine survival, as this pleiotropic gene is known to modulate cysteine protease activity and stability (Rausch and Hastings, 2015). In addition, there was an indication of higher selection pressure on a *lyst-1* orthologue, a regulator of endosomal trafficking in *C. elegans* polarized epithelial cells (de Souza et al., 2007), that may share the same function in *H. contortus* and thus contribute to efficient processing of protein material from the intestinal lumen. This suggests that regulation of the proteolytic pathways in vaccine survivors may result in improved survival. While the precise function of cysteine peptidases is hard to infer *in silico*, current knowledge from *in vitro* studies points to their role in the proteolytic cascade responsible for degrading haemoglobin or immunoglobulin G (Williamson et al., 2003). Perhaps worms that over-express these proteins may either maintain blood coagulation and digestion, or are able to degrade host IgG stimulated by the vaccine challenge (Munn et al., 1997; Ekoja and Smith, 2010) to evade the vaccine response, or some combination of both. Indeed the vaccine is proposed to disrupt digestion in the worm gut by blocking the function of the intestinal proteases it targets. Processing of ingested proteins by an alternative proteolytic pathway may improve the survival and/or fecundity of worms suffering dietary restriction. In addition, the over-expression of a myo-inositol-1 phosphate synthase in vaccine survivors may also support this theory as this gene is known to act on lipid storage (Ashrafi et al., 2003) and in the defecation cycle (Tokuoka et al., 2008), both critical in the digestion process, and hence impacting worm growth and lifespan. Interestingly, the most highly differentially expressed genes show a low level of expression in worms from the control group, suggesting that the vaccine response may have induced their overexpression in the vaccine survivors or alternatively, that the vaccine selects for natural variation in expression of these genes. Additional transcriptomic evaluation of the offspring of each worm subpopulation, before and after vaccine exposure, would help confirm this observation and distinguish between a regulatory response to vaccine-induced immunity and genetic differences influencing gene expression.

Whilst this study focuses on a species of veterinary significance, our findings may have relevance to other species. Indeed our results suggest that *H. contortus* may be able to compensate for vaccine-mediated immunity after vaccine exposure and a similar situation may apply in other parasitic nematode systems.

## Conclusions

Our data suggest that parasite populations surviving Barbervax^®^ immunisation are able to optimize their proteolytic machinery, involving both peptidases and regulators of lysosome trafficking, and display better lipid storage and/or defecation abilities which may enhance survival in the face of a robust vaccine-induced immune response. While our experiment was not designed to detect genetic selection to the vaccine response, an “evolve and resequencing” approach to contrast changes in allele frequencies in vaccinated and unvaccinated populations through time, across multiple generations of vaccine challenge, could help resolve the potential for adaptation following vaccination.

## Acknowledgements

We thank Stephen Doyle for advice and comments on the manuscript and the biological services staff at MRI for their expert animal care. JAC, NH, AM, AT and MB are supported by the Wellcome Trust via their core funding of the Wellcome Trust Sanger Institute (grant 206194). JAC, NH, AT, MB, KM, RL, ED and CB are supported by BBSRC grant BB/M003949/1 (BUG), GS has received the support of the EU in the framework of the Marie-Curie FP7 COFUND People Programme, through the award of an AgreenSkills (grant agreement n° 267196) and AgreenSkills+ fellowships (grant agreement n°609398). The funders had no role in study design, data collection and analysis, decision to publish, or preparation of the manuscript.

## Supporting information

**Supplementary Figure1. Principal component analysis (PCA) of transcript counts measured in worms collected in vaccinated or control sheep**

PCA is a dimensionality reduction method that makes use of transcript counts to define a new set of unrelated components. Coordinates of every pool of worms considered for analysis is plotted against first two components and correlate with similarities between pools. The first PCA axis explains 36% of total variance and relates to differences between the two considered experimental groups, *i.e.* worms exposed to the vaccine response (V) or the control group (C).

**Supplementary Figure2. Number of differentially expressed genes found by each of the two implemented methods**

Total number of significantly differentially expressed genes found by at least one of the two methods (DESeq2, VOOM, or both (intersecting)) are plotted according to their regulation pattern, *i.e* up or down-regulated in the vaccine survivors, to their estimated fold change, i.e. log2FC > 2, 1 or 0.

**Supplementary Table 1.** Faecal egg count and worm volumes recovered at necropsy and RNA-seq library details

**Supplementary Table 2.** List of primer sequences used for qPCR validation

**Supplementary Table 3.** Complete list of differentially expressed genes

**Supplementary Table 4.** Transcript count of the six most differentially expressed genes in pools of male worms from each sheep

**Supplementary Table 5.** Significantly enriched G0 terms associated with the differentially expressed genes

